# Diffusive and advective fluid flow shapes chemoautotrophic bacterial communities and sulfur mineralogy in hydrothermal sediments off Milos

**DOI:** 10.1101/2025.09.02.673406

**Authors:** Joely Marie Maak, Clemens Röttgen, Birte Winkelhues, Eirini Anagnostou, Wolfgang Bach, Solveig I. Bühring, Andrea Koschinsky, Jianlin Liao, Christoph Vogt, Enno Schefuß, Marcus Elvert

## Abstract

Hydrothermal fluid flow not only shapes mineral deposition on the ocean floor but also creates ecological niches by altering temperature and energy availability. In these niches, microbial life thrives and has an additional, often unrecognized impact on mineral formation. In a newly discovered vent field in medium depths off Milos, Greece, we show how contrasting hydrothermal regimes host fundamentally different bacterial metabolisms. Diffusive flow fosters acidic, sulfate-rich conditions that promote kaolinization and pyrite formation. Fatty acid δ^13^C values down to −39‰ indicate acetyl-CoA-based sulfate reduction as the main metabolism, likely contributing to pyrite formation. In contrast, vigorous venting delivers hot, acidic, carbon-rich, and sulfate-depleted fluids that sustain the activity of sulfide-oxidizing chemoautotrophs. Fatty acid δ^13^C values of up to −4‰ and elemental sulfur accumulation provide evidence of their activity. Without such hydrothermal influence, little microbial activity and only quartz-rich sediments can be observed. Combined multivariate lipid analyses highlight Eh as the strongest environmental control, with increasing average chain length of fatty acids as an adaptation to (hydro-)thermal stress. These results illustrate how different types of fluid flow influence the activity of chemoautotrophic bacterial communities that are involved in the formation of characteristic sulfur minerals in hydrothermal environments.

## Introduction

Submarine hydrothermal vents are hotspots of life, where fluid-rock interactions shape microbial activity. These systems emerge in tectonically or volcanically active regions, such as ocean floor spreading zones or volcanic flanks (Tarasov et al. 2005) and can broadly be categorized into deep-sea and shallow-water hydrothermal systems mostly defined by depth and by light availability. Due to high hydrostatic pressure, deep-sea systems can reach temperatures of 400°C and above (Tarasov et al. 2005, Koschinsky et al. 2008, Baker et al. 2012, Reeves et al. 2014), while shallow systems exhibit lower temperatures (∼100 to 200°C; Caramanna et al. 2021, Khimasia et al. 2021). Especially systems that occur in shallower depths in the photic zone (≤ 200 m) are dynamic natural laboratories for studying how microorganisms adapt to extreme conditions as they are more accessible than their deep counterparts. Generally, shallow hydrothermal systems are characterized by gas emission of mostly CO_2_, H_2_S, and CH_4_ (Kleint et al. 2019, Caramanna et al. 2021). Mixing of emanating fluids with seawater causes cooling of the fluids to temperatures suitable for life and creates steep geochemical gradients, fostering communities of specialized chemolithoautotrophic bacteria and archaea (Le Moine Bauer et al. 2023) and leading to the precipitation of minerals (Lubetkin et al. 2018).

The Aegean Sea off Greece is known for several areas with submarine fluid and gas venting associated with the Hellenic Volcanic Arc. One of such tectonically active areas lies off the coast of Milos Island (Dando et al. 1995). Although the last volcanic eruption occurred around 90 ka ago, residual heat from the underlying magma system continues to fuel hydrothermal venting activity (Dando et al. 1995, Zhou et al. 2021). Previous reports document venting up to 110 meters water depth covering an area of about 35 km^2^ with temperatures of up to 122.4°C (Dando et al. 1995, Khimasia et al. 2020). The gases emitted from these shallow vents are predominantly composed of CO_2_ (often exceeding 90%), with smaller contributions of H_2_S, CH_4_, and H_2_ (Dando et al. 1995, Dando et al. 2000). The vent fluids in these areas are sulfide-rich, exhibit elevated concentrations of arsenic and mercury, and are acidic with pH values down to 5 (Price et al. 2013, Roberts et al. 2021). Near the shore, yellow-orange arsenic and sulfur-rich or brown manganese- and iron-rich mineral precipitates form when the fluids mix with seawater (Godelitsas et al. 2015). Microbial communities are dominated by Campylobacteria (formerly Epsilonproteobacteria, Waite et al. 2017, mostly *Arcobacteraceae, Sulfurimonadaceae*, and *Sulfurovaceae*) and Gammaproteobacteria (e.g., Giovannelli et al. 2013, Le Moine Bauer et al. 2023).

Campylobacteria can play a significant role in sulfur, nitrogen, and hydrogen cycling at hydrothermal systems (e.g., Campbell et al. 2006). These chemoautotrophic microorganisms utilize different pathways for carbon assimilation, including the Calvin-Benson-Bassham (CBB) cycle, but also the reductive tricarboxylic acid (rTCA) cycle, both of which have been found to be abundant in the shallow regions off Milos (Callac et al. 2017). The rTCA cycle is often used by microorganisms in energy-limited environments like hydrothermal systems (e.g., Campbell and Cary 2004, Maak et al. 2025). Another possible pathway includes the acetyl-CoA pathway, which has been observed at deep-sea hydrothermal systems (Nakagawa and Takai 2008). These pathways produce characteristic stable carbon isotope (δ^13^C) values in the corresponding biomass, including polar membrane-derived fatty acids; organisms employing the rTCA cycle generate the most positive δ^13^C values and hence lowest carbon isotopic fractionation (ε) relative to the CO_2_ substrate (ε_rTCA_ = −2 to −13‰), while the other two produce larger fractionations (ε_acetyl-CoA_ = −20 to −36‰, ε_CBB_ = −10 to −22‰, Preuß et al. 1989, Hayes 2001, House et al. 2003, Reeves et al. 2014).

During our recent research cruise M192 in August 2023, we observed the occurrence of hydrothermal venting down to depths of 210 meters and with fluid temperatures of up to 180°C (Nomikou et al., under review). In contrast to the shallower regions, the deeper areas also feature chimney structures with varying elemental compositions from which the microbial communities have not yet been examined. Here, we provide first insights into the newly discovered vent fields in deeper regions off Milos and combine temperature, redox potential (Eh), pH, dissolved inorganic carbon (DIC), δ^13^C DIC, dissolved organic carbon (DOC), as well as the most important anion (sulfate, chloride, bromide) and element concentrations (Al, Fe, Si, Ca) in the porewater with the existing mineralogy and the variability of fatty acids (concentration and δ^13^C values). Fatty acids are produced as intermediate products and storage lipids by microorganisms (e.g., Alvarez et al. 1997), but could also be degradation products from intact polar lipids (e.g., Schouten et al. 2010). Fatty acids are exclusively produced by bacteria and eukaryotes, with bacteria being the most likely producers of fatty acids in our setting, as they are known to be important in carbon and sulfur cycling in hydrothermal systems (e.g., Cao et al. 2014). Since hydrothermal fluid flow not only generates thermal and chemical gradients that shape microbial metabolisms but also drives mineral formation and alteration, integrating fatty acid data with porewater geochemistry and mineralogy is essential to understand where biological and geological processes intersect. We specifically examined three sediment cores from different depths, representing: (i) a “Background” core devoid of hydrothermal influence, (ii) a pronounced redox gradient in Eh and pH values (“Gradient” core), and (iii) fully reduced conditions reaching temperatures of up 100°C at the bottom (“Hot vent” core).

### Experimental Procedures

During the METEOR research cruise M192 from 08. August until 05. September 2023, the entire vent system around Milos was examined in detail (see Cruise Report, Bühring et al. 2023). Two cores at hydrothermal-impacted sites (Gradient: GeoB25519−13, 163 m water depth; Hot vent: GeoB25526−4, 108 m water depth), and one Background core (GeoB25514−13, 103 m water depth) were sampled using a multicorer (MUC, Background and Gradient core) or the remotely operated vehicle (ROV) MARUM-SQUID (Hot vent core, Nowald et al. 2016). The push core taken with the ROV could be retrieved in the desired location in the vicinity of active venting, while cores taken with a MUC were not camera guided, thus verifying hydrothermal influence in these cores was done on deck after recovery. Since the Hot vent core was taken using a pushcore of a smaller diameter, less porewater could be recovered and this core was not measured for DOC and major element concentrations (details see below).

Temperatures in the sediment were measured with a temperature stick using the MARUM ROV SQUID a few centimeters next to the actual location of the Hot vent core. Measured temperatures were 45°C at 5 cm below seafloor (cmbsf) and 160°C at 30 cmbsf. For the topmost centimeter of the sediment, we used 15°C obtained from the overlying bottom water. To obtain a downcore temperature profile, a thermal diffusion model was applied to the in-situ measured temperatures using input parameters for diffusivity of 3 × 10^−7^ m^2^ s^−1^ for sandy sediments and flow velocity of 0.021 m h^−1^ (see Sollich et al. 2017). The data was interpolated in RStudio (v. 4.0.3; R Core Team, 2020) using the *approxfun* function provided in the vegan package (v. 2.5−6; Oksanen et al. 2020), with method set to linear interpolation in 100 µm steps.

Sediment cores were sampled for porewater using Rhizones (0.15 µm pore width) in 2 cm sample resolution, sliced in one cm intervals and stored at −20°C until further analyses. Porewaters were prepared for various shipboard and shore-based analyses using the following procedures: pH and Eh values were measured immediately (WTW pH electrode SenTix® 940 for pH and WTW SenTix® ORP-T 900 with a Ag/AgCl electrode for Eh), for measurements of DIC and DOC samples were stored at +4°C, for ion chromatography (IC), samples were filtered through 0.2 μm polyethersulfone (PES) filters under a laminar flow hood; for inductively coupled plasma optical emission spectroscopy (ICP-OES), samples were filtered and acidified to pH < 2 using suprapure HCl, and then stored in acid-cleaned LDPE bottles. Measurements of DIC and DOC (measured as non-purgeable organic carbon) were performed using an Analytik Jena multiN/C 2100s at MARUM, University of Bremen. For analyses of DIC δ^13^C values porewater aliquotes of 1 ml were transferred into exetainer glass vials, pre-flushed with CO_2_ free air and acidified with 100 µl of 45% phosphoric acid, and let react overnight. Headspace CO_2_ was then analyzed in single measurements at MARUM using a Delta Ray Isotope ratio Infrared Spectrometer from Thermo Fisher Scientific coupled to an ASX−7100 Autosampler from CETAC.

Major anions (Cl^-^, SO_4_^2-^, Br^-^) in the porewater were measured on 150-fold diluted samples at Constructor University Bremen using a Metrohm 761 Compact IC system, equipped with a Metrosep A Supp 5−150/4.0 column. The eluent was a mixture of sodium carbonate (Na_2_CO_3_) and sodium bicarbonate (NaHCO_3_) in deionized water at final concentrations of 1.0 mM and 3.2 mM, respectively. Analytical precision and accuracy were evaluated using IAPSO standard seawater, yielding an uncertainty of less than 2% for the targeted anions.

Total element concentration (Al, Fe, Si, Ca) in the porewater was analyzed at MARUM, University of Bremen, using Inductively Coupled Plasma Optical Emission Spectroscopy (ICP-OES) on a Varian Vista Pro instrument. The system was fitted with an argon gas humidifier, seaspray nebulizer, and a cyclonic spray chamber. Samples were diluted 20-fold prior to analysis. Calibration was performed using a matrix-matched mixture of laboratory single-element standards across five concentration levels. Data quality was validated through repeated measurements of IAPSO P-series standard seawater (salinity 34.994), with external precision better than 5.5% and external accuracy better than 1.5%. Internal precision was consistently better than 3%. Due to dilution requirements, the detection limits for less abundant elements were: Al (3.7 μM) and Fe (0.7 μM), all measured with internal precision better than 3%.

X-ray diffraction (XRD) pattern analysis of freeze-dried and ground sediments was performed using a Bruker D8 Discover Diffractometer, equipped with a Cu-tube (k_α_ 1.541 Å, 40 kV, 40 mA), a fixed divergence slit of 0.25°, and a 90 samples changer and a monochromatization via energy discrimination on the highest resolution Linxeye detector system. The scan range was 3 to 65 2theta, with a step size of 0.016° 2theta and a scan step time of 0.7 seconds. Elemental compositions of sedimentary grains were further examined using a Bruker energy dispersive X-ray (EDX) spectrometer with a XFlash 6|30 detector coupled to a Zeiss SUPRA 40 field emission gun scanning electron microscope (SEM). Averaged relative standard deviations were better than 2% for most measurements, except for oxygen and iron, which could be up to 6% in cases of high abundance of the individual element (>50 at.%). Samples were carbon-coated prior to SEM work.

Total sulfur (TS) and total organic carbon (TOC) were measured using a LECO CS 744. Portions of 100 mg powdered and homogenized sediment samples were dissolved in excess 12.5% hydrochloric acid (HCl) to remove carbonates. The TOC and TS content was analyzed in single measurements by subsequent combustion of the decalcified samples and consecutive identification of SO_2_ and CO_2_ in non-dispersive infrared (NDIR) cells. Concentrations were determined via multi-point regression calibration using certified LECO reference material.

For fatty acid analyses, about 10 grams of sliced sediment samples (every cm) were freeze-dried and extracted following a modified Bligh and Dyer protocol (Sturt et al. 2004). For later quantification, an internal standard (2Me-C_18:0_) of known concentration was added to the samples. To obtain fatty acids, 50% of the TLE was used for saponification, followed by the formation of fatty acid methyl esters (FAMEs) using boron trifluoride in methanol (Elvert et al. 2003). FAMEs were identified by gas chromatography coupled to mass spectrometry (GC-MS) using an Agilent 7820 GC equipped with a 30 m J&W DB−5MS column (0,25 mm internal diameter (i.d), 0,25 µm film thickness) and interfaced to an Agilent 5077E mass-selective 200 detector. Quantification of FAMEs was performed by a FOCUS GC using a flame ionization detector (GC-FID, Thermo Fisher Scientific) equipped with a 30 m Restek Rxi−5ms column (0.25 mm i.d., 0.25 µm film thickness). The injector was adjusted at 260°C with the oven temperature initially being held constant at 70°C for 2 minutes, then increased to 150°C over the next 4 minutes, raised to 320°C over the next 42 minutes and held at this temperature for 20 minutes, with nitrogen gas (N_2_) as carrier gas and a constant flow rate of 1.3 ml/min. Concentration of FAMEs was calculated based on single measurements in relation to the internal standard 2Me-C_18:0_. Contents are displayed in µg g^−1^ dried sediment. Average chain length (ACL) was calculated from the concentrations and carbon chain lengths of all detected fatty acids that would fall in the bacterial range (C_12_ to C_20_).

Compound specific δ^13^C values of FAMEs were determined using a Thermo Fisher Scientific Trace GC equipped with a TriPlus autosampler and a J&W DB−1 GC column (30 m length, 0.25 mm i.d., 0.5 µm film thickness) and coupled to a Conflo II to a Thermo Finnigan MAT 252 isotope mass spectrometer. Injection was performed in splitless mode. The temperature program of the GC-system started with initially 120°C, 3 min isothermal, was then subsequently raised to the final temperature of 320°C at a rate of 5°C/min, 22 min isothermal. Helium was used as the carrier gas at a flow rate of 1.2 ml/min. All δ^13^C values are expressed in the per mil scale (‰) relative to Vienna Pee Dee Belemnite (VPDB) and are referenced to an in-house laboratory CO_2_ reference gas with an analytical error of <0.3 ‰, and were corrected for the addition of carbon from methanol during derivatization. As monounsaturated fatty acids were eluting very closely to each other, they were combined in the evaluation of their δ^13^C values.

A non-metric multidimensional scaling (NMDS) analysis was performed on the fatty acid distribution. The ordination was performed on single compounds using the Bray-Curtis distance measure. Fatty acid data was log-transformed but not scaled (to reduce the influence of outliers, Ramette 2007), while environmental variables were scaled to a mean of 0 and unit variance using the *decostand* function from the vegan package (v. 2.5−6; Oksanen et al. 2020). Post-correlation of environmental variables, based on 999 permutations yielded significant to highly significant levels with p ≤ 0.05. In addition, pairwise Spearman rank correlations were calculated between all environmental variables and individual fatty acid compounds. For each pair, the correlation coefficient (ρ) and associated p-value were determined using the *cor*.*test* function in R. P-values were corrected for multiple testing using the Benjamini–Hochberg method (Benjamini and Hochberg 1995). All fatty acid–environment pairs with an adjusted p-value (padj) ≤ 0.05 and spearmans ρ ≥0.6 or ≤−0.6 were retained (results see supplementary Tab. 1). At the Hot vent site, estimates for pH, Eh, Cl^-^, and SO_4_ ^2-^ at 0.5 cmbsf. were made due to missing data, for which the respective values from 2.5 cmbsf. were used, as these parameters show little variability with depth.

## Results

### Porewater measurements

The three sediment cores show clear differences in the porewater parameters. The Background core shows only minimal, if any, changes in composition relative to seawater (main parameters: pH: 7.4±0.06, Eh: 180±14, Cl^-^: 609±17 mM, SO_4_^2-^: 31±0.8 mM, Fig. 2, supplementary Fig. 1). In contrast, the Gradient core shows continuous changes in Eh and pH (changes from 197 to −107 in Eh and from 7.6 to 5.9 in pH) towards the base of the core, whereas Cl^-^ and SO_4_^2-^ are constantly similar to the Background with values of 593±12 mM and 30±0.6 mM, respectively (Fig. 2, supplementary Fig. 1). The Hot vent core is reducing and acidic throughout the entire core length (pH: 5.8±0.2, Eh: −256±2.2, Cl^-^: 501±12 mM, SO_4_^2-^: 17±1.2 mM, Fig. 2). DIC concentrations in the Hot vent core range from 4.2 to 8.4 mM while the Background and Gradient cores show lower concentrations (mean of 2.4±0.3 mM). DIC δ^13^C values range from −3.0‰ to +16.9‰ in the entire data set with up to +16.9‰ detected in the Hot vent core, continuously more negative values around 0‰ in the Background core, and most negative values of −3.0‰ found at the topmost sample in the Gradient core (Fig. 3). The dominant major element concentrations of Al, Si, Fe, and Ca measured only for the Background and Gradient cores were relatively similar, with the exception of substantially higher Al and Fe contents in the Gradient (Al: up to 0.56 mg/L, Fe: up to 3.7 mg/L) than in the Background core (Al: below 0.12 mg/L, Fe: below 0.9 mg/L, supplementary Fig. 2).

### Mineralogy, total sulfur, and total carbon

TOC contents were generally highest in the Background (0.43 to 0.55%) and show a slight downcore decrease in the Gradient core (0.28 to 0.19%, Fig. 3). In the Hot vent core, remnants of seagrass lead to an additional peak in TOC at 9 to 17 cm sediment depth (up to 0.60%). Sulfur is below 1% in the Background, but present throughout the entire Gradient core (between 6 to 16%, Fig. 3). At the Hot vent site, sulfur peaks in the topmost centimeter with about 6%, followed by a strong decrease to about 1% (3 to 17 cm). The mineralogical assemblage, measured by XRD, shows that the Background and the Hot vent core are similar with high contents of quartz (50 to 80%, Fig. 2) and relatively high contents of feldspar (up to 20%). Both cores differ in the amount of phyllosilicate present (Background below 20%, Hot vent core up to 35%). The Hot vent core is the only one containing alunite (up to 3%), while Al_2_Si_2_O_5_(OH)_4_ polymorphs are missing. Sulfides make up <1% in the Background and Hot vent core. The Gradient core shows strong differences compared to the other two cores with elevated contents in phyllosilicate (up to 43%) and sulfides (up to 28%). Al_2_Si_2_O_5_(OH)_4_ polymorphs are abundant throughout (17±7%). Indeed, in the Gradient core, SEM imaging revealed abundant kaolinite and discrete cubic pyrite crystals at every depth horizon (Fig. 4); both phases notably increasing in grain size with depth. The sediment grains of the topmost core sample displayed a continuous coating with Fe(oxy)hydroxide (nearly pure Fe by EDX, Fig. 4, supplementary Tab. 2) that was not detected by XRD and is hence suspected to be amorphous. Kaolinite is the most abundant Al_2_Si_2_O_5_(OH)_4_ polymorph (up to 32%), but nacrite and halloysite were also detected by XRD and can make <5% of the sediments in some intervals. In the Hot vent core, SEM–EDX, similar to XRD, showed predominantly quartz fragments and some siliceous microfossils. Dispersed elemental sulfur micrograins (≈6 wt.% by XRD, Figs. 2 and 4) coat quartz and siliceous microfossil surfaces, documenting ongoing sulfide oxidation at the ocean floor interface as observed in the shallow vents (Gilhooly et al. 2014). Titanium-bearing oxide particles (∼19 at%, likely anatase, see XRD results), formed on top of quartz grains, were present in the Hot vent core in varying abundances.

### Fatty acid contents and δ^13^C values

Bacterial fatty acid contents (C_12_ to C_20_) are highest in all the uppermost layers (Fig. 3), with contents of 10.0 µg/g (Background), 15.6 µg/g (Gradient), and 7.2 µg/g (Hot vent). In total, contents decrease to 2.9 µg/g (Background), 1.3 µg/g (Gradient), and 0.8 µg/g (Hot vent). Monounsaturated fatty acids are most prominent in the Gradient core, with 7.3 µg/g in the top layer, decreasing to 0.2 µg/g at 22 cm depth. The Background shows contents of 3.9 µg/g of monounsaturated fatty acids at the top that is decreasing to about 0.2 µg/g in the lower sediment layers. At the Hot vent site, contents decrease from 2.8 µg/g to 0.1 µg/g. The fatty acid C_16:0_ is dominating in all cores and at all depths. All sediment cores show a wide variety in δ^13^C values, ranging between −31.9 (C_16:0_) and −23.2‰ (C_28:0_) in the Background, between −39.1 (10-Me-C_16_) to −23.9‰ (C_28:0_) in the Gradient, and between −32.2 (C_30:0_) and −4‰ (C_16:1_) in the Hot vent, which shows the highest diversity of all cores. Overall, some fatty acids in the Hot vent are enriched in ^13^C, mostly C_16:1_ (up to −4‰), C_17:1_ (up to −14.4‰), and C_18:1_ (up to −11.3‰), while the Gradient core shows fatty acids with strong ^13^C-depletion (down to −39.1‰, see supplementary Fig. 1). Including the DIC δ^13^C and the resulting fractionation (ε) in fatty acids, we observe fractionation between −29.8‰ to −21.3‰ in the Background, between −39.6‰ to −21.0‰ in the Gradient core, and between −44.6‰ to −11.6‰ in the Hot vent core (Fig. 3). Notably, the largest apparent fractionations in the Hot vent occur in the lowermost sediments, which likely do not reflect active carbon assimilation or new biomass production.

## Interpretation and Discussion

### Separation of sedimentary regimes into advective and diffusive fluid flow

Distinguishing between (1) focused, advective discharge and (2) dispersed or diffusive fluid flow in hydrothermal sedimentary regimes (e.g., Price et al. 2007, Pop Ristova et al. 2017) is critical for redox zonation, mineral precipitation, and microbial activity. At Milos hydrothermal vents, constant sulfate and chloride values in the Gradient core as opposed to the change observed in pH and Eh (Fig. 1, supplementary Fig. 1), suggest that this environment is dominated by slow and diffusive flow of hydrothermal fluids. Here, fluids with low-redox potential and mildly acidic pH rise slowly from below, combined with elevated Fe and Al levels. This allows seawater to penetrate relatively deep into the sedimentary regime, as suggested by the constantly elevated sulfate concentrations (Fig. 2b). At the Hot vent, strongly advective and directed fluid flow causes uniform, extreme conditions of negative Eh and low pH associated with a substantial depletion in sulfate (Fig. 2c). Seawater and, therefore oxygen, only penetrates a few millimeters deep into the sediments. The increasingly elevated DIC concentrations and ^13^C-enriched values of DIC from the Background to the Gradient to the Hot vent core underscore the increasing influence of rising magmatic hydrothermal fluids (Figs. 2 and 3). The phenomenon of strong enrichment in ^13^C in the DIC in hydrothermally influenced sediments has also been observed in the near-shore shallow systems with values of up to +12‰ (Callac et al. 2017). This isotopic effect cannot (solely) be the result of degassing of isotopically light ^12^CO_2_, as potential solubility of CO_2_ would be ∼28.6 mM at atmospheric pressure and ambient air temperature (∼1 bar and 30°C, Duan and Sun 2003) as opposed to the maximum of 10 mM we observe in our cores (Fig. 2). Furthermore, no outgassing was observed on deck after sediment core retrieval. One possible explanation is that CO_2_ gas rising from depth dissolves progressively into cooler water near the seafloor, and the isotope fractionation during its conversion to bicarbonate enriches the dissolved carbon in ^13^C. In the Background core, constant sulfate, chloride, DIC, pH, and Eh values prove absence of any fluid flow.

**Figure 1.**
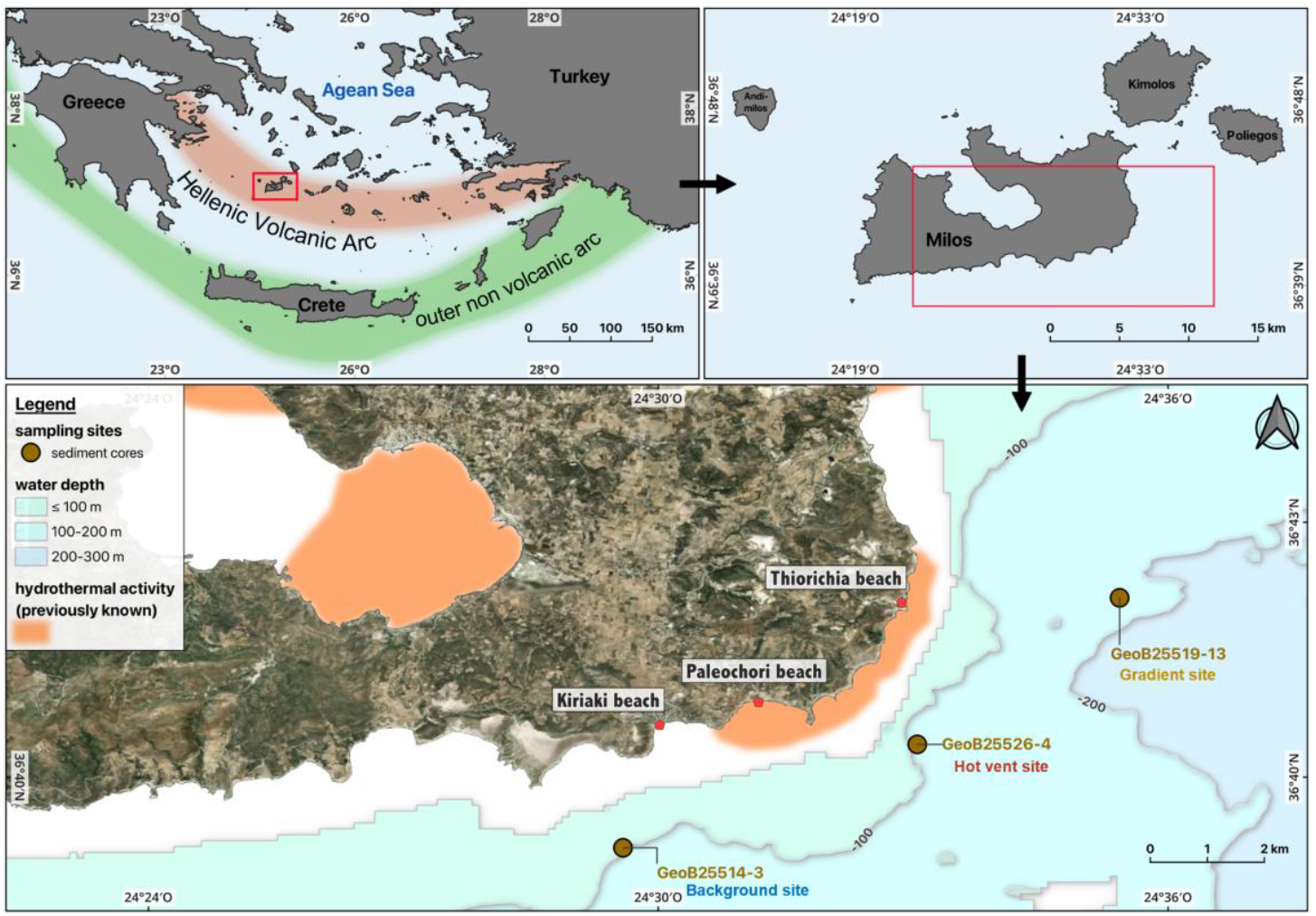
Location of the study area near the Greek island of Milos within the Hellenic volcanic arc (top left and right) as well as the previously known areas of hydrothermal activity and the sampling sites of the sediment cores used in this study (bottom). The satellite image is from © Apple Maps (2024). Bathymetry contour lines were extracted from the 125 m resolution multibeam grid from the M192 cruise (Nomikou et al., under review).

**Figure 2.**
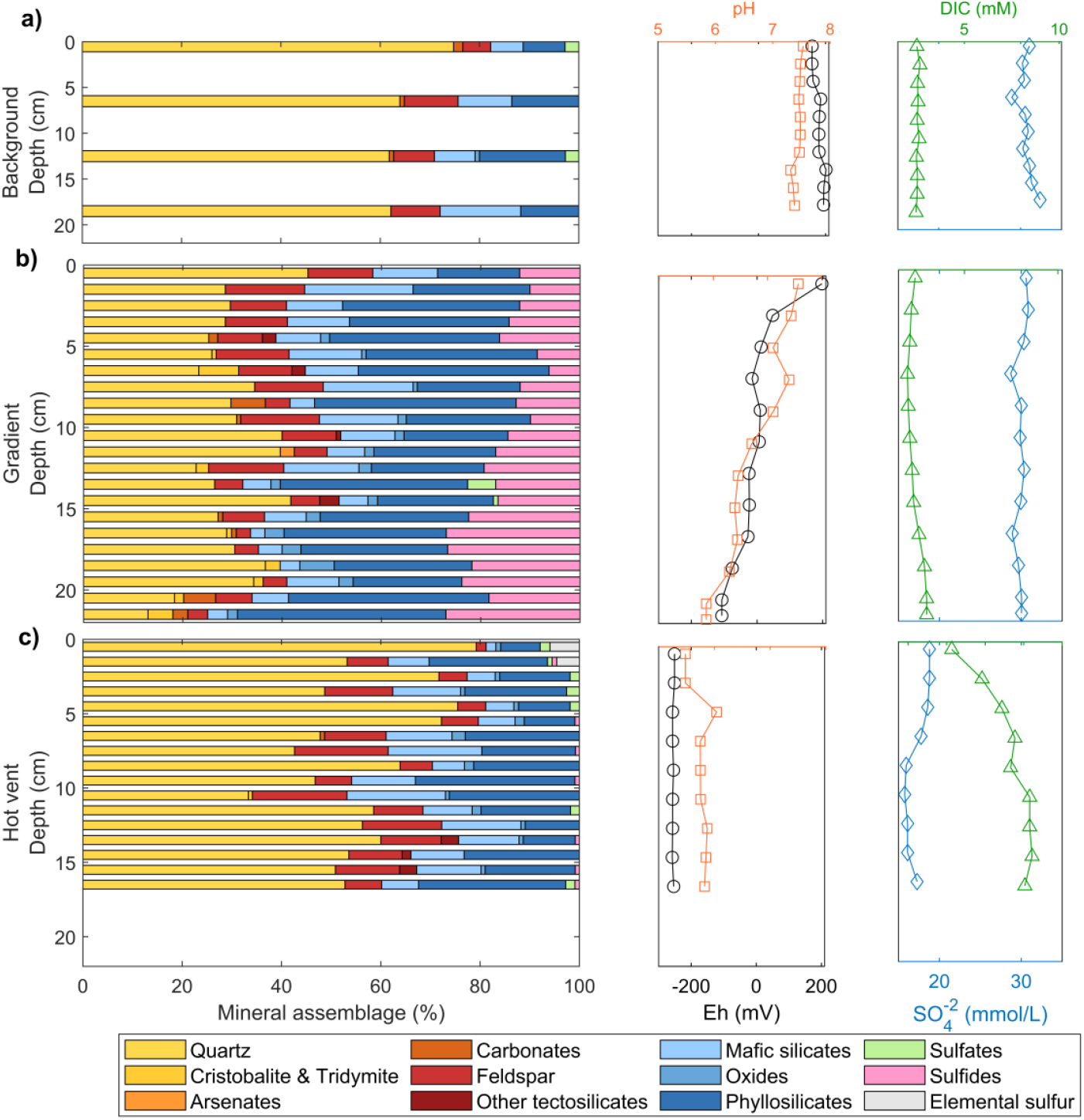
Mineralogy and porewater profiles from the three cores: a) Background; b) Gradient; and c) Hot vent. Crystalline mineral assemblages are shown as stacked horizontal bars (% of total). Sulfide phases and elemental sulfur appear only in the hydrothermally influenced cores. Porewater profiles include redox potential (Eh in mV; black circles), pH (orange squares), sulfate (SO_4_^2-^ in mM; blue diamonds), and dissolved inorganic carbon (DIC in mM; green triangles).

**Figure 3.**
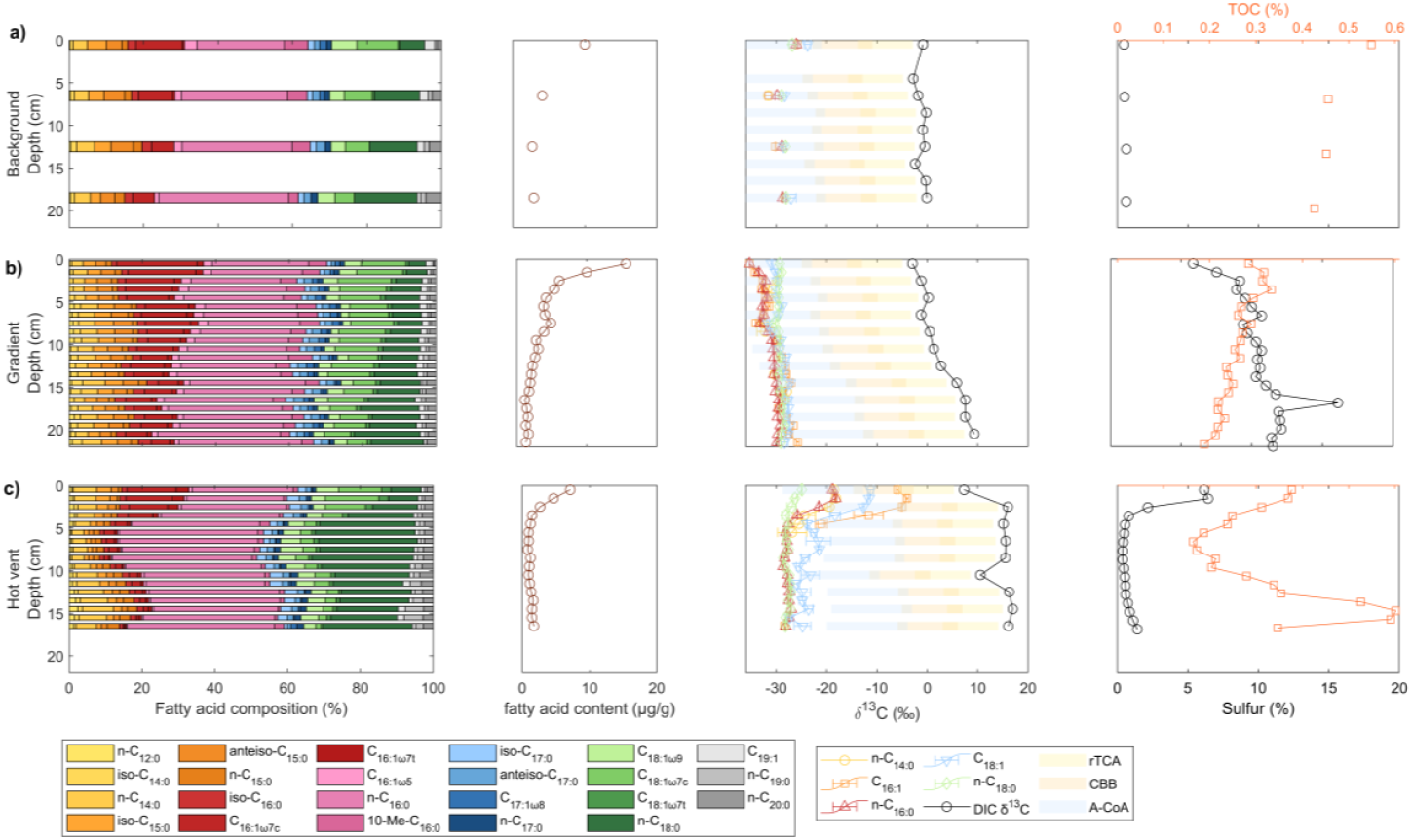
Bacterial fatty acid composition (in %), content (in µg/g) and compound-specific δ^13^C values of specific fatty acids and δ^13^C values of DIC (in ‰), as well as TOC and total sulfur content (both in %) in a) Background, b) Gradient, and c) Hot vent core. Stacked horizontal bars show the relative abundance of individual fatty acids C_12_–C_20_ each in a distinct color. δ^13^C values of selected fatty acids (C_14:0_, C_16:1_, C_16:0_, C_18:1_, C_18:0_) with error bars represent duplicate measurements. Typical fractionation ranges of chemoautotrophy are displayed for reductive tricarboxylic acid cycle (rTCA, ε = −2 to −13‰, shaded in yellow), the Calvin-Benson-Bassham cycle (CBB ε = −10 to −22‰, shaded in orange) and the reductive acetyl CoA pathway (A-CoA, ε = −20 to −36‰, shaded in blue; Hayes 2001, House et al. 2003, Reeves et al. 2014). Total sulfur includes all phases of sulfur (sulfates, sulfides, and elemental sulfur).

### Impact of fluid flow type on mineralogy in sedimentary hydrothermal regimes

Hydrothermal fluids, both advective and diffusive, deliver heat, reduced metals, sulfur, silica, and various trace elements into the upper sediment layers (e.g., Bortnikov et al. 2014, Hsu et al. 2024). As these fluids encounter cooler, oxidizing porewaters and detrital sediments, they precipitate mineral phases that both record past fluid–rock interactions and shape modern biogeochemical cycling. The extent of fluid-mineral reactions depends on permeability, fluid geochemistry, and temperature (Buatier et al. 1995, Fulignati 2020). In our study, focusing on the newly discovered hydrothermal sediments from deeper regions off Milos, abundant clay, pyrite, marcasite, and elemental sulfur confirm active hydrothermal alteration and sulfide deposition in fluid flow-affected sediments (Fig. 2b). Indeed, hydrothermal alteration of feldspar and mica minerals to kaolinite-group minerals and the precipitation of pyrite and marcasite is common in hydrothermal systems, with marcasite often formed under acidic conditions (Murowchick and Barnes 1986, Meunier 1995, Njoya et al. 2006). Similarly, in the very shallow regions off Milos, Callac et al. (2017) also reported elemental sulfur together with pyrite and marcasite only in white-capped, hydrothermally influenced sediments. In the Gradient core, these features are very pronounced, and likely a result of elongated fluid-rock interactions due to a slow and diffusive flow of fluids. On top of the Gradient core is an oxidated iron-rich layer (Fig. 4a, not visible by XRD), suggesting recent or poorly crystalline Fe-oxyhydroxide formation in the topmost section of the sediment that is affected by oxygen penetration. In contrast, the Hot vent sediment core is located in a zone with strong advective flow, closer to a heat source. Despite the clear signs of hydrothermal discharge in porewater, the sediments show much less evidence for hydrothermal alteration manifested in the complete lack of kaolinite and sulfide, which commonly make up 50% of the Gradient core. The less pronounced hydrothermal alteration of the Hot vent sediments suggests much shorter durations of fluid-sediment interaction. However, some hydrothermal precipitates are present, including minimal amounts of pyrite, elemental sulfur, alunite, and TiO_2_ (see Fig. 2c and Fig. 4d-f). Elemental sulfur is abundant only in the top two centimeters and is dispersed as micron-scale grains in the void space between other mineral grains but alunite is a minor constituent of the sediments throughout. Both phases could be related to the discharge of magmatic SO_2_, which can disproportionate to form elemental sulfur and sulfuric acid. The latter can transform feldspar and white mica to alunite and quartz (Hemley et al. 1969). The sulfate depletion means that the circulating seawater must have precipitated anhydrite upon heating to temperatures above 140°C (Bischoff and Seyfried 1978). The enrichment of DIC in the porewater (Fig. 2c) shows magmatic volatile influx and the low pH recorded for the porewater is testament of high contents of dissolved CO_2_. The abundance of authigenic TiO_2_ coating other mineral grains indicates some transport of Ti which could also point to acidic conditions affected by magmatic emanations. From the described hydrothermal mineral phases both sulfides and elemental sulfur could be the product of biological activity. While SO_2_-disproportionation can produce elemental sulfur, it could also be the product of sulfide oxidation to elemental sulfur (e.g., Cron et al. 2019). Since elemental sulfur exclusively occurs in the top section of the sediments at the Hot vent, this would be consistent with bacterial H_2_S oxidation rather than disproportionation of SO_2_. Pyrite formation in the Gradient core could be microbially mediated as well (e.g., Thiel et al. 2019, Jia et al. 2023).

**Figure 4.**
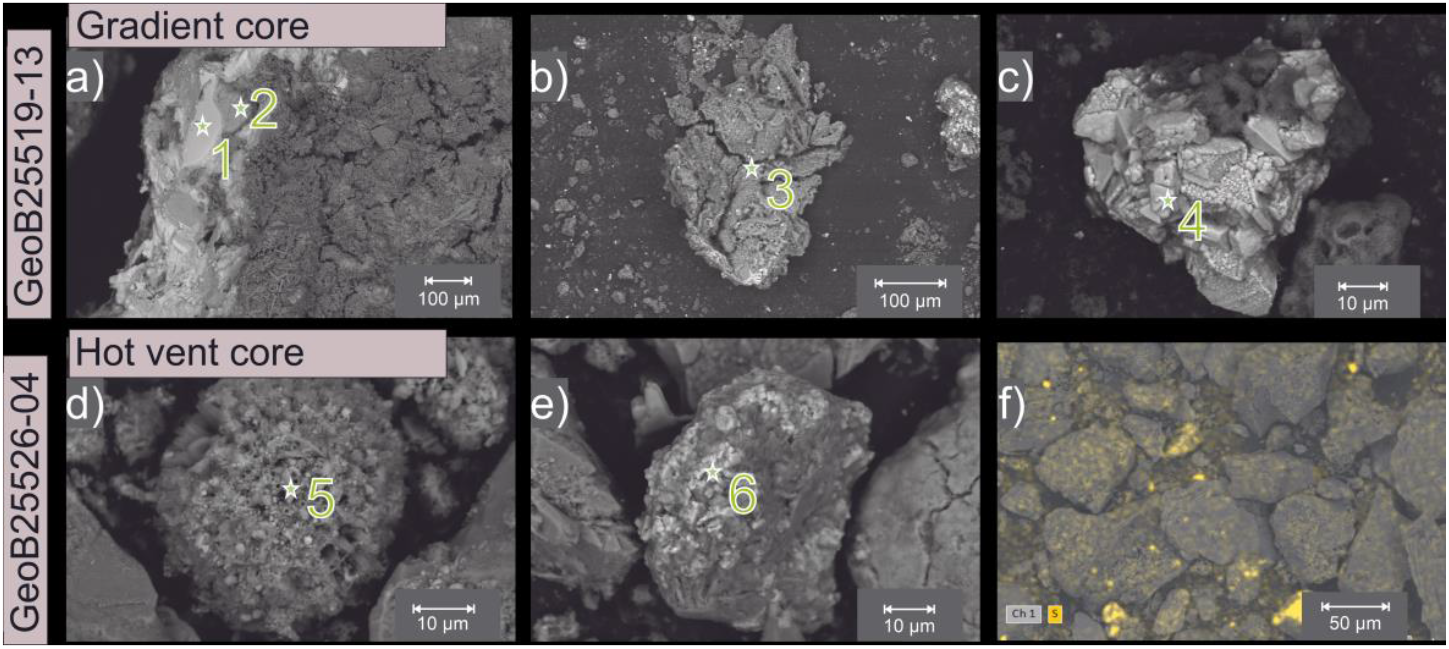
SEM–EDX micrographs of key mineral phases in Gradient (a–c) and Hot-vent (d–f) cores. (a) Fe-crust atop the Gradient core, cut by bright pyrite veins; the surrounding gray matrix is composed of almost pure iron precipitate (likely amorphous), iron oxides, and Al- and Si-oxides. (b) Kaolinite fragments with ∼Si/Al/O proportions (∼14:12:74 at.%), characteristic of Al-rich clay. (c) Cubic pyrite crystal. (d) Sphere dominated by SiO_2_ (∼90 at.% Si+O) with minor contents of Fe, Al and S, consistent with amorphous silica precipitates mingled with hydrothermal sulfide/oxide phases. (e) Ti-bearing grains enriched in Ti relative to Si (∼19:4 at.%), most likely a TiO_2_ polymorph (rutile or anatase). (f) Overview of Hot vent sediment (1–2 cm) showing dispersed micron-sized grains of elemental sulfur (according to XRD ≈ 6 wt.%). EDX results of point measurements (1 to 6) are shown in supplementary Tab. 2.

### Biological control on element cycling and sulfur mineral formation in hydrothermal sediments

Non-metric multidimensional scaling (NMDS) of fatty acid distributions clearly separates the Background, Gradient, and Hot vent samples into distinct clusters, indicating that each regime hosts a unique bacterial fatty acid composition (Fig. 5). Environmental vectors fitted onto the NMDS ordination reveal that Eh and pH are the strongest drivers of fatty acid variability (R^2^ = 0.80 and 0.67, p ≤ 0.001), followed by DIC concentration (R^2^ = 0.62, p ≤ 0.001), the concentrations of the major anions Cl^-^, Br^-^, and SO_4_^2-^ (all R^2^ ≈ 0.58–0.60, p ≤ 0.001), temperature (R^2^ = 0.54, p ≤ 0.001), and ACL (R^2^ = 0.45, p ≤ 0.001; Fig. 5). The isotopic composition of DIC has a strong correlation to lipid clustering as well (R^2^ = 0.79, p ≤ 0.001; Fig. 5), which is likely caused by the strong correlation between DIC concentration and its isotopic value (see Figs. 2 and 3). Generally, the dominant control of DIC and SO_4_^2-^ concentrations suggest chemoautotrophic sulfur-metabolizing microorganisms as most active in all the hydrothermally influenced sedimentary regimes. Sulfur isotope studies have already suggested that microorganisms are involved in the sulfur cycle in the shallow hydrothermal systems off Milos (Gilhooly et al. 2014, Houghton et al. 2019).

**Figure 5.**
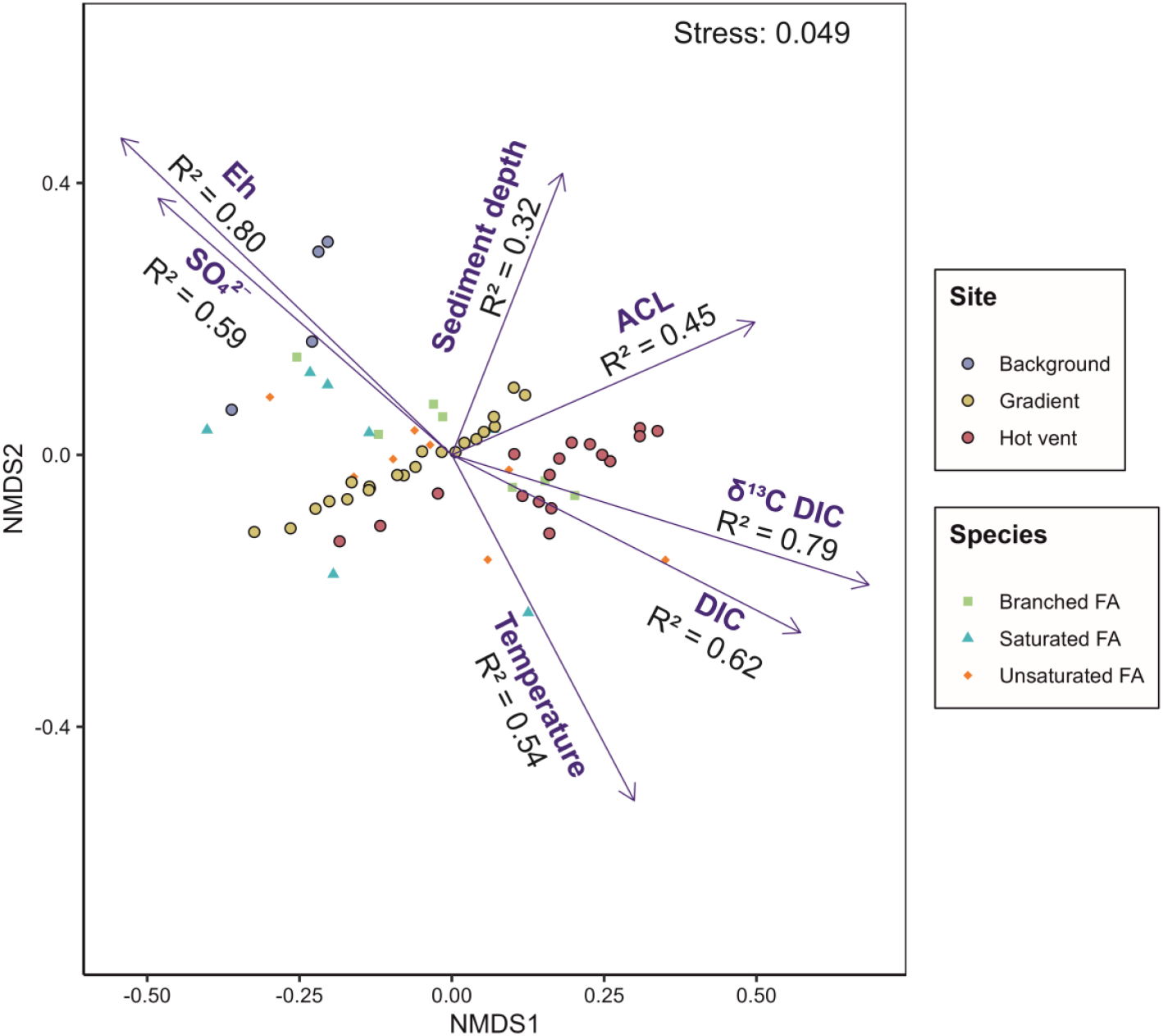
Non-metric multidimensional scaling (NMDS) biplot showing the dissimilarity of the different hydrothermal regimes based on the bacterial fatty acid distribution (C_12_ to C_20_). The proportion of variation explained by each environmental factor is provided as R^2^ values, the individual p-values are displayed in Tab. 1. ACL: average chain length.

On the other hand, temperature is less correlated in our case and is not the main driver of metabolic activity as found in a previous study on lipid diversity close to the beach (Sollich et al. 2017). In the deeper regions, temperature was not shown to be the main parameter controlling bacterial fatty acid variability. Similar to the NMDS, the correlation analyses using Spearmans ρ revealed that Eh is the dominant control, with highest Spearmans ρ values found for Eh with *iso*-C_17:0_ (ρ = 0.94), n-C_15:0_, n-C_17:0_, *anteiso*-C_15:0_, C_16:1ω7c_, and C_16:1ω5_ fatty acids (all ρ > 0.82, p (adjusted using Benjamini and Hochberg 1995) ≪ 10^−10^, supplementary Tab. 1). Most of these fatty acids are also highly abundant in the cores (Fig. 3) and similarly controlled by the pH value, which co-varies strongly with Eh (supplementary Tab. 1, Fig. 2b and c). Meanwhile, temperature did not show correlations with individual fatty acids but did show a strong relationship with the ACL (including bacterial fatty acids of C_12_ to C_20_) in the Hot vent core until 61°C (R^2^ = 0.95, p ≪ 0.001, supplementary Fig. 1) which is likely close to the upper limit for bacterial life in this environment. The observed correlation between temperature and ACL likely reflects an adaptation to extreme environmental conditions. This trend is also apparent in the NMDS, where ACL increases along a similar axis as sediment depth (Fig. 5), suggesting a combined influence of high temperature, low pH, and low Eh. Microbial lipid compositions often mirror adaptive responses of the source organisms to changing conditions – such as exposure to toxic elements (Ghosh et al. 2018) or elevated temperatures (Brassell et al. 1986, Sollich et al. 2017, Garcia et al. 2024), by adjusting membrane lipid structure to regulate ion permeability (Boyd et al. 2011, Garcia et al. 2024). Such adaptations help maintain the electrochemical gradients across membranes that are essential for energy conservation processes, including ATP synthesis, making them critical for efficient cellular bioenergetics (Valentine and Valentine 2004). The drop in abundance of bacteria above 60°C further correlates with the temperature ceilings for metagenomic functional diversity (Ruhl et al. 2022) and operational taxonomic units (Sharp et al. 2014) measured in geothermal springs in Canada and New Zealand.

The Hot vent sediment with low pH and Eh, high temperatures, and high DIC concentrations creates an environment where only thermophilic and acidotolerant chemoautotrophic microbes are able to thrive (e.g., Deng et al. 2023). Bacteria in such extreme environments often include sulfide oxidizers that use the rTCA cycle (Inagaki et al. 2004, Takai et al. 2006, Mino et al. 2014, Giovannelli et al. 2016, Maak et al. 2025) and produce elemental sulfur as an intermediate during the oxidation of sulfide with oxygen or nitrate at the water–sediment interface (e.g., Fuseler and Cypionka 1995, Cron et al. 2019, Wang et al. 2022). The rTCA cycle fractionates less and produces ^13^C-enriched monounsaturated C_16_ and C_18_ fatty acids (Maak et al. 2025). We found high abundances of these fatty acids with compound-specific δ^13^C values more positive relative to layers below (only depleted by −11.6‰ relative to DIC at +7.3‰, Fig. 3) alongside elevated elemental sulfur contents in the upper centimeters of the Hot vent sediment (Fig. 3c). However, the importance of rTCA in hydrothermal sediments at Milos is obviously limited to those vent systems with extremely strong advective fluid flow from deeper depths, as typical δ^13^C values were not observed in the Gradient core as well as in coastal sediments of shallower regions (Callac et al. 2017).

In total, concentrations of fatty acids are significantly higher in the Gradient core than in the Background or Hot vent cores, indicating an *in-situ* production of bacterial fatty acids. Typically, biomass produced by other carbon assimilation cycles such as the CBB would fall in the range of −10 to −22‰ (Hayes 2001, House et al. 2003). However, in the Gradient core, fatty acids show a much stronger depletion of up to −37.0‰ in C_16:1_ and C_18:1_ relative to DIC, and 10-Me-C_16_ (a fatty acid often produced by sulfate reducers, e.g., Dowling et al. 1986) even shows a ^13^C depletion of up to −39.6‰, which is not consistent with the activity of the CBB cycle. A possible pathway that would result in a stronger carbon isotope fractionation than the CBB and rTCA cycle would be the reductive acetyl-CoA pathway (Hayes 2001, Fig. 3). This pathway is restricted to anaerobic chemoautotrophic microbes, is often connected to sulfate-reducing bacteria (Ragsdale 2004, Nakagawa and Takai 2008), and can cause very high ε values as shown for other sulfate reducers like *Desulfotomaculum acetoxidans* (ε_FA-DIC_ ≈ −31‰; Londry and Des Marais 2003). Previous studies have shown that diffusive venting in hydrothermal systems can cause elevated activity of sulfate-reducing bacteria (Jørgensen et al. 1990, Roerdink et al. 2024). Indeed, other studies on these shallow systems have already shown high rates of sulfate reduction that have been observed at and below 40°C and pH between 5 and 7, which decline at increasing pCO_2_ (Bayraktarov et al. 2013). Moreover, in shallow areas around Milos, sulfate-reducing genera (e.g., Gomez-Saez et al. 2017), as well as functional genes of sulfate reducers (*dsrA*) have been reported (Callac et al. 2017, Le Moine Bauer et al. 2023). It is therefore very plausible that the Gradient site is influenced by bacterial sulfate-reduction associated with a highly fractionating chemoautotrophic pathway, such as the acetyl-CoA pathway.

## Conclusions

Our findings underscore the importance of the presence and the intensity of fluid flow characteristics on shaping the biogeochemistry, mineral assemblages, and bacterial life in sedimentary hydrothermal regimes. In particular, we show that even moderate changes in redox and pH conditions can cause distinct mineralogy and support functionally different microbial communities, each adapted to their specific geochemical niche. The diffusive Gradient core with slow migration of low-temperature fluids favors hydrothermal kaolinization and pyrite formation and fosters a microbial community dominated by sulfate-reducing bacteria utilizing the acetyl-CoA cycle, that might contribute to sulfide mineral precipitation. In contrast, the mineral phases in the Hot vent core seem to be hydrothermally altered to lesser extents while the geochemical and fatty acid distribution is strongly affected by the rapid advective flow of acidic fluids. Bacterial life is limited to the topmost centimeters, where sulfide-oxidizers use the rTCA cycle and lead to an increase in elemental sulfur deposition and dissolved sulfate concentrations. This integrated geochemical, mineralogical, and lipid biomarker approach offers new insights into biogeochemical niches at newly discovered medium to shallow hydrothermal systems and underlines the importance of fluid flow dynamics in structuring subsurface mineralogy as well as microbial life.

## Data availability

The supplementary data supporting the findings of this study are available as a separate file and in the PANGAEA database (data has been sent to PANGAEA but has not been published yet).

## Acknowledgements

We thank Detlef Korte and the crew of RV METEOR for their support during cruise M192. We thank the German Research Vessel Fleet Coordination Centre for granting ship time and providing logistics. We also extend thanks to the team of the ROV SQUID (N. Nowald, T.Leymann, R. Rehage, S.Schillai) for operating the underwater vehicle and their help in sample recovery. This study was supported by the Deutsche Forschungsgemeinschaft (DFG, German Research Foundation) under Germany’s Excellence Strategy (EXC−2077—390741603) and an independent cruise proposal no GPF 20−2/019. We also acknowledge financial support for open-access publishing from the University of Bremen’s Open Access Publication Fund.

## Ethics Statement

This study did not involve research on human participants or vertebrate animals. All prevailing local, national, and international regulations and conventions, as well as normal scientific ethical practices, including those related to the Convention on Biological Diversity and the Nagoya Protocol, were respected.

## Author Contributions

Samples were collected by JMM, CR, ES, JL, WB, SIB, AK, and ME. JMM, CR, ES, and ME designed the study, and JMM, CR, BW, EA, JL, and CV carried it out. JMM, CR, and BW interpreted the data. JMM prepared the manuscript with contributions from all co-authors.

---

**Table 1.** Environmental parameters from the non-metric multidimensional scaling (NMDS) with a p value <0.05 (statistically significant). Eh, temperature, DIC, δ^13^C DIC, SO_4_^2-^, sediment depth, and average chain length (ACL) are depicted in Fig. 5.

|  | NMDS1 | NMDS2 | R<sup>2</sup> | p-value |
| --- | --- | --- | --- | --- |
| **Eh** | −0.68 | 0.58 | 0.80 | 0.001 |
| **δ<sup>13</sup>C DIC** | 0.86 | −0.24 | 0.79 | 0.001 |
| **pH** | −0.73 | 0.36 | 0.67 | 0.001 |
| **DIC** | 0.72 | −0.33 | 0.62 | 0.001 |
| **Cl<sup>−</sup>** | −0.59 | 0.51 | 0.60 | 0.001 |
| **SO<sub>4</sub><sup>2−</sup>** | −0.60 | 0.47 | 0.59 | 0.001 |
| **Br<sup>−</sup>** | −0.61 | 0.46 | 0.58 | 0.001 |
| **Temperature** | 0.37 | −0.64 | 0.54 | 0.001 |
| **ACL** | 0.62 | 0.24 | 0.45 | 0.001 |
| **Sediment**<br>**depth (cm)** | 0.23 | 0.52 | 0.32 | 0.001 |
| **Si** | 0.11 | 0.51 | 0.27 | 0.005 |
| **DOC** | 0.37 | 0.20 | 0.18 | 0.018 |
| **Fe** | −0.08 | −0.40 | 0.17 | 0.022 |

## References

Alvarez, H. M., Pucci, O. H. & Steinbüchel, A. (1997) Lipid storage compounds in marine bacteria. Appl. Microbiol. Biotechnol., 47. 10.1007/s002530050901

Baker, E. T., Walker, S. L., Embley, R. W. & de Ronde, C. E. J. (2012) High-resolution hydrothermal mapping of Brothers Caldera, Kermadec Arc. Econ. Geol., 107. 10.2113/econgeo.107.8.1583

Bayraktarov, E., Price, R. E., Ferdelman, T. G. & Finster, K. (2013) The pH and pCO2 dependence of sulfate reduction in shallow-sea hydrothermal CO2 – venting sediments (Milos Island, Greece). Front. Microbiol., Volume 4 - 2013. 10.3389/fmicb.2013.00111

Benjamini, Y. & Hochberg, Y. (1995) Controlling the false discovery rate: a practical and powerful approach to multiple testing. J. R. Stat. Soc., Ser. B, Stat. Methodol., 57. 10.1111/j.2517-6161.1995.tb02031.x

Bischoff, J. L. & Seyfried, W. E. (1978) Hydrothermal chemistry of seawater from 25°C to 350°C. Am. J. Sci., 278. 10.2475/ajs.278.6.838

Bortnikov, N. S., Simonov, V. A., Amplieva, E. E. & Borovikov, A. A. (2014) Anomalously high concentrations of metals in fluid of the Semenov modern hydrothermal system (Mid-Atlantic Ridge, 13°31′ N): LA-ICP-MS study of fluid inclusions in minerals. Dokl. Earth Sci., 456. 10.1134/S1028334X14060221

Boyd, E. S., et al. (2011) Temperature and pH controls on glycerol dibiphytanyl glycerol tetraether lipid composition in the hyperthermophilic crenarchaeon Acidilobus sulfurireducens. Extremophiles, 15. 10.1007/s00792-010-0339-y

Brassell, S. C., Eglinton, G., Marlowe, I. T., Pflaumann, U. & Sarnthein, M. (1986) Molecular stratigraphy: a new tool for climatic assessment. Nature, 320. 10.1038/320129a0

Buatier, M. D., Früh-Green, G. L. & Karpoff, A. M. (1995) Mechanisms of Mg-phyllosilicate formation in a hydrothermal system at a sedimented ridge (Middle Valley, Juan de Fuca). Contrib. Mineral. Petrol., 122. 10.1007/s004100050117

Bühring, S. I., et al. (2023) Bridging hydrothermal sites along the Hellenic Arc off Milos from shallow to deep, Cruise No. M192/1 and M192/2, August 08 - September 05, 2023, Piraeus - Limassol. METEOR-Berichte. 10.48433/cr_m192

Callac, N., et al. (2017) Modes of carbon fixation in an arsenic and CO2-rich shallow hydrothermal ecosystem. Sci. Rep., 7. 10.1038/s41598-017-13910-2

Campbell, B. J. & Cary, S. C. (2004) Abundance of reverse tricarboxylic acid cycle genes in free-living microorganisms at deep-sea hydrothermal vents. Appl. Environ. Microbiol., 70. 10.1128/AEM.70.10.6282-6289.2004

Campbell, B. J., Engel, A. S., Porter, M. L. & Takai, K. (2006) The versatile ε-proteobacteria: key players in sulphidic habitats. Nat. Rev. Microbiol., 4. 10.1038/nrmicro1414

Cao, H., et al. (2014) Microbial sulfur cycle in two hydrothermal chimneys on the southwest Indian Ridge. mBio, 5. 10.1128/mbio.00980-13

Caramanna, G., Sievert, S. M. & Bühring, S. I. (2021) Submarine shallow-water fluid emissions and their geomicrobiological imprint: a global overview. Front. Mar. Sci., 8. 10.3389/fmars.2021.727199

Cron, B., Henri, P., Chan, C. S., Macalady, J. L. & Cosmidis, J. (2019) Elemental sulfur formation by Sulfuricurvum kujiense is mediated by extracellular organic compounds. Front. Microbiol., 10. 10.3389/fmicb.2019.02710

Dando, P. R., et al. (2000) Hydrothermal studies in the Aegean Sea. Phys. Chem. Earth, Part B: Hydrology, Oceans and Atmosphere, 25. 10.1016/S1464-1909(99)00112-4

Dando, P. R., et al. (1995) Gas venting rates from submarine hydrothermal areas around the island of Milos, Hellenic Volcanic Arc. Cont. Shelf Res., 15. 10.1016/0278-4343(95)80002-U

Deng, W., et al. (2023) Strategies of chemolithoautotrophs adapting to high temperature and extremely acidic conditions in a shallow hydrothermal ecosystem. Microbiome, 11. 10.1186/s40168-023-01712-w

Dowling, N. J. E., Widdel, F. & White, D. C. (1986) Phospholipid ester-linked fatty acid biomarkers of acetate-oxidizing sulphate-reducers and other sulphide-forming bacteria. Microbiology, 132. 10.1099/00221287-132-7-1815

Duan, Z. & Sun, R. (2003) An improved model calculating CO2 solubility in pure water and aqueous NaCl solutions from 273 to 533 K and from 0 to 2000 bar. Chem. Geol., 193. 10.1016/S0009-2541(02)00263-2

Elvert, M., Boetius, A., Knittel, K. & Jørgensen, B. B. (2003) Characterization of specific membrane fatty acids as chemotaxonomic markers for sulfate-reducing bacteria involved in anaerobic oxidation of methane. Geomicrobiol. J., 20. 10.1080/01490450303894

Fulignati, P. (2020) Clay minerals in hydrothermal systems. Minerals, 10. 10.3390/min10100919

Fuseler, K. & Cypionka, H. (1995) Elemental sulfur as an intermediate of sulfide oxidation with oxygen byDesulfobulbus propionicus. Arch. Microbiol., 164. 10.1007/BF02525315

Garcia, A. A., Chadwick, G. L., Liu, X.-L. & Welander, P. V. (2024) Identification of two archaeal GDGT lipid–modifying proteins reveals diverse microbes capable of GMGT biosynthesis and modification. PNAS, 121. 10.1073/pnas.2318761121

Ghosh, D., Bhadury, P. & Routh, J. (2018) Coping with arsenic stress: adaptations of arsenite-oxidizing bacterial membrane lipids to increasing arsenic levels. Microbiologyopen, 7. 10.1002/mbo3.594

Gilhooly, W. P., et al. (2014) Sulfur and oxygen isotope insights into sulfur cycling in shallow-sea hydrothermal vents, Milos, Greece. Geochem. Trans., 15. 10.1186/s12932-014-0012-y

Giovannelli, D., et al. (2016) Sulfurovum riftiae sp. nov., a mesophilic, thiosulfate-oxidizing, nitrate-reducing chemolithoautotrophic epsilonproteobacterium isolated from the tube of the deep-sea hydrothermal vent polychaete Riftia pachyptila. Int. J. Microbiol., 66. 10.1099/ijsem.0.001106

Giovannelli, D., d’Errico, G., Manini, E., Yakimov, M. & Vetriani, C. (2013) Diversity and phylogenetic analyses of bacteria from a shallow-water hydrothermal vent in Milos island (Greece). Front. Microbiol., 4. 10.3389/fmicb.2013.00184

Godelitsas, A., et al. (2015) Amorphous As-sulfide precipitates from the shallow-water hydrothermal vents off Milos Island (Greece). Mar. Chem., 177. 10.1016/j.marchem.2015.09.004

Gomez-Saez, G. V., et al. (2017) Relative importance of chemoautotrophy for primary production in a light exposed marine shallow hydrothermal system. Front. Microbiol., 8. 10.3389/fmicb.2017.00702

Hayes, J. (2001) Fractionation of carbon and hydrogen isotopes in biosynthetic processes. Rev. Mineral. Geochem., 43. 10.2138/gsrmg.43.1.225

Hemley, J. J., Hostetler, P. B., Gude, A. J. & Mountjoy, W. T. (1969) Some stability relations of alunite. Econ. Geol., 64. 10.2113/gsecongeo.64.6.599

Houghton, J. L., et al. (2019) Spatially and temporally variable sulfur cycling in shallow-sea hydrothermal vents, Milos, Greece. Mar. Chem., 208. 10.1016/j.marchem.2018.11.002

House, C. H., Schopf, J. W. & Stetter, K. O. (2003) Carbon isotopic fractionation by Archaeans and other thermophilic prokaryotes. Org. Geochem., 34. 10.1016/S0146-6380(02)00237-1

Hsu, F.-H., et al. (2024) Geochemical indications of hydrothermal fluid through sediments within the Geolin Mounds and Mienhua Volcano hydrothermal fields, southernmost Okinawa Trough. Deep-Sea Res. I: Oceanogr. Res. Pap., 207. 10.1016/j.dsr.2024.104293

Inagaki, F., Takai, K., Nealson, K. H. & Horikoshi, K. (2004) Sulfurovum lithotrophicum gen. nov., sp. nov., a novel sulfur-oxidizing chemolithoautotroph within the ε-Proteobacteria isolated from Okinawa Trough hydrothermal sediments. Int. J. Syst. Evol. Microbiol., 54. 10.1099/ijs.0.03042-0

Jia, F., et al. (2023) Sulphate-reducing bacteria-mediated pyrite formation in the Dachang Tongkeng tin polymetallic deposit, Guangxi, China. Sci. Rep., 13. 10.1038/s41598-023-38827-x

Jørgensen, B. B., Zawacki, L. X. & Jannasch, H. W. (1990) Thermophilic bacterial sulfate reduction in deep-sea sediments at the Guaymas Basin hydrothermal vent site (Gulf of California). Deep-Sea Res. I: Oceanogr. Res. Pap., 37. 10.1016/0198-0149(90)90099-H

Khimasia, A., Alessio, R. & and Pichler, T. (2020) Hydrothermal areas, microbial mats and sea grass in Paleochori Bay, Milos, Greece. Int. J. Maps, 16. 10.1080/17445647.2020.1748131

Khimasia, A., Renshaw, C. E., Price, R. E. & Pichler, T. (2021) Hydrothermal flux and porewater geochemistry in Paleochori Bay, Milos, Greece. Chem. Geol., 571. 10.1016/j.chemgeo.2021.120188

Kleint, C., et al. (2019) Geochemical characterization of highly diverse hydrothermal fluids from volcanic vent systems of the Kermadec intraoceanic arc. Chem. Geol., 10.1016/j.chemgeo.2019.119289

Koschinsky, A., et al. (2008) Hydrothermal venting at pressure-temperature conditions above the critical point of seawater, 5°S on the Mid-Atlantic Ridge. Geology, 36. 10.1130/g24726a.1

Le Moine Bauer, S., et al. (2023) Structure and metabolic potential of the prokaryotic communities from the hydrothermal system of Paleochori Bay, Milos, Greece. Front. Microbiol., 13. 10.3389/fmicb.2022.1060168

Londry, K. L. & Des Marais, D. J. (2003) Stable carbon isotope fractionation by sulfate-reducing bacteria. Appl. Environ. Microbiol., 69. 10.1128/AEM.69.5.2942-2949.2003

Lubetkin, M., et al. (2018) Nontronite-bearing tubular hydrothermal deposits from a Galapagos seamount. Deep-Sea Res. II: Top. Stud. Oceanogr., 150. 10.1016/j.dsr2.2017.09.017

Maak, J. M., et al. (2025) The energy-efficient reductive tricarboxylic acid cycle drives carbon uptake and transfer to higher trophic levels within the Kueishantao shallow-water hydrothermal system. Biogeosciences, 22. 10.5194/bg-22-1853-2025

Meunier, A. 1995. Hydrothermal alteration by veins. In: Velde, B. (ed.) Origin and Mineralogy of Clays: Clays and the Environment. Berlin, Heidelberg: Springer Berlin Heidelberg. 978-3-662-12648-6

Mino, S., et al. (2014) Sulfurovum aggregans sp. nov., a hydrogen-oxidizing, thiosulfate-reducing chemolithoautotroph within the Epsilonproteobacteria isolated from a deep-sea hydrothermal vent chimney, and an emended description of the genus Sulfurovum. Int. J. Syst. Evol. Microbiol., 64. 10.1099/ijs.0.065094-0

Murowchick, J. B. & Barnes, H. L. (1986) Marcasite precipitation from hydrothermal solutions. Geochim. Cosmochim. Acta, 50. 10.1016/0016-7037(86)90214-0

Nakagawa, S. & Takai, K. (2008) Deep-sea vent chemoautotrophs: diversity, biochemistry and ecological significance. FEMS Microbiol. Ecol., 65. 10.1111/j.1574-6941.2008.00502.x

Njoya, A., et al. (2006) Genesis of Mayouom kaolin deposit (western Cameroon). Appl. Clay Sci., 32. 10.1016/j.clay.2005.11.005

Nowald, N., Ratmeyer, V. & Wefer, G. MARUM-Squid - a powerful, yet compact 2000 m ROV system designed for marine research operations from smaller vessels. OCEANS 2016 MTS/IEEE Monterey. 10.1109/OCEANS.2016.7761353

Oksanen, J., et al. (2020) vegan community ecology package version 2.5-7 November 2020. 10.32614/CRAN.package.vegan

Pop Ristova, P., Pichler, T., Friedrich, M. W. & Bühring, S. I. (2017) Bacterial diversity and biogeochemistry of two marine shallow-water hydrothermal systems off Dominica (Lesser Antilles). Front. Microbiol., 8. 10.3389/fmicb.2017.02400

Preuß, A., Schauder, R., Fuchs, G. & Stichler, W. (1989) Carbon isotope fractionation by autotrophic bacteria with three different CO2 fixation pathways. Z. Naturforsch. C., 44. 10.1515/znc-1989-5-610

Price, R. E., Amend, J. P. & Pichler, T. (2007) Enhanced geochemical gradients in a marine shallow-water hydrothermal system: Unusual arsenic speciation in horizontal and vertical pore water profiles. Appl. Geochem., 22. 10.1016/j.apgeochem.2007.06.010

Price, R. E., et al. (2013) Processes influencing extreme As enrichment in shallow-sea hydrothermal fluids of Milos Island, Greece. Chem. Geol., 348. 10.1016/j.chemgeo.2012.06.007

Ragsdale, S. W. (2004) Life with carbon monoxide. Crit. Rev. Biochem. Mol. Biol., 39. 10.1080/10409230490496577

Ramette, A. (2007) Multivariate analyses in microbial ecology. FEMS Microbiol. Ecol., 62. 10.1111/j.1574-6941.2007.00375.x

Reeves, E. P., et al. (2014) Microbial lipids reveal carbon assimilation patterns on hydrothermal sulfide chimneys. Environ. Microbiol., 16. 10.1111/1462-2920.12525

Roberts, H., Price, R., Brombach, C.-C. & Pichler, T. (2021) Mercury in the hydrothermal fluids and gases in Paleochori Bay, Milos, Greece. Mar. Chem., 233. 10.1016/j.marchem.2021.103984

Roerdink, D. L., et al. (2024) Hydrothermal activity fuels microbial sulfate reduction in deep and distal marine settings along the Arctic Mid Ocean Ridges. Front. Mar. Sci., 10. 10.3389/fmars.2023.1320655

Ruhl, I. A., et al. (2022) Microbial functional diversity correlates with species diversity along a temperature gradient. mSystems, 7. 10.1128/msystems.00991-21

Schouten, S., Middelburg, J. J., Hopmans, E. C. & Sinninghe Damsté, J. S. (2010) Fossilization and degradation of intact polar lipids in deep subsurface sediments: a theoretical approach. Geochim. Cosmochim. Acta, 74. 10.1016/j.gca.2010.03.029

Sharp, C. E., et al. (2014) Humboldt’s spa: microbial diversity is controlled by temperature in geothermal environments. ISME J., 8. 10.1038/ismej.2013.237

Sollich, M., et al. (2017) Heat stress dictates microbial lipid composition along a thermal gradient in marine sediments. Front. Microbiol., 8. 10.3389/fmicb.2017.01550

Sturt, H. F., Summons, R. E., Smith, K., Elvert, M. & Hinrichs, K.-U. (2004) Intact polar membrane lipids in prokaryotes and sediments deciphered by high-performance liquid chromatography/electrospray ionization multistage mass spectrometry—new biomarkers for biogeochemistry and microbial ecology. Rapid Commun. Mass Spectrom., 18. 10.1002/rcm.1378

Takai, K., et al. (2006) Sulfurimonas paralvinellae sp. nov., a novel mesophilic, hydrogen- and sulfur-oxidizing chemolithoautotroph within the Epsilonproteobacteria isolated from a deep-sea hydrothermal vent polychaete nest, reclassification of Thiomicrospira denitrificans as Sulfurimonas denitrificans comb. nov. and emended description of the genus Sulfurimonas. Int. J. Syst. Evol. Microbiol., 56. 10.1099/ijs.0.64255-0

Tarasov, V. G., Gebruk, A. V., Mironov, A. N. & Moskalev, L. I. (2005) Deep-sea and shallow-water hydrothermal vent communities: two different phenomena? Chem. Geol., 224. 10.1016/j.chemgeo.2005.07.021

Thiel, J., Byrne, J. M., Kappler, A., Schink, B. & Pester, M. (2019) Pyrite formation from FeS and H2S is mediated through microbial redox activity. Proc. Natl. Acad. Sci., 116. 10.1073/pnas.1814412116

Valentine, R. C. & Valentine, D. L. (2004) Omega-3 fatty acids in cellular membranes: a unified concept. Prog. Lipid Res., 43. 10.1016/j.plipres.2004.05.004

Waite, D. W., et al. (2017) Comparative genomic analysis of the class Epsilonproteobacteria and proposed reclassification to Epsilonbacteraeota (phyl. nov.). Front. Microbiol., 8. 10.3389/fmicb.2017.00682

Wang, T., et al. (2022) The pathway of sulfide oxidation to octasulfur globules in the cytoplasm of aerobic bacteria. Appl. Environ. Microbiol., 88. 10.1128/aem.01941-21

Zhou, X., Kuiper, K., Wijbrans, J., Boehm, K. & Vroon, P. (2021) Eruptive history and ^40^Ar/^39^Ar geochronology of the Milos volcanic field, Greece. GChron., 3. 10.5194/gchron-3-273-2021

