## supplementary data for "Diffusive and advective fluid flow shapes chemoautotrophic bacterial communities and sulfur mineralogy in hydrothermal sediments off Milos"

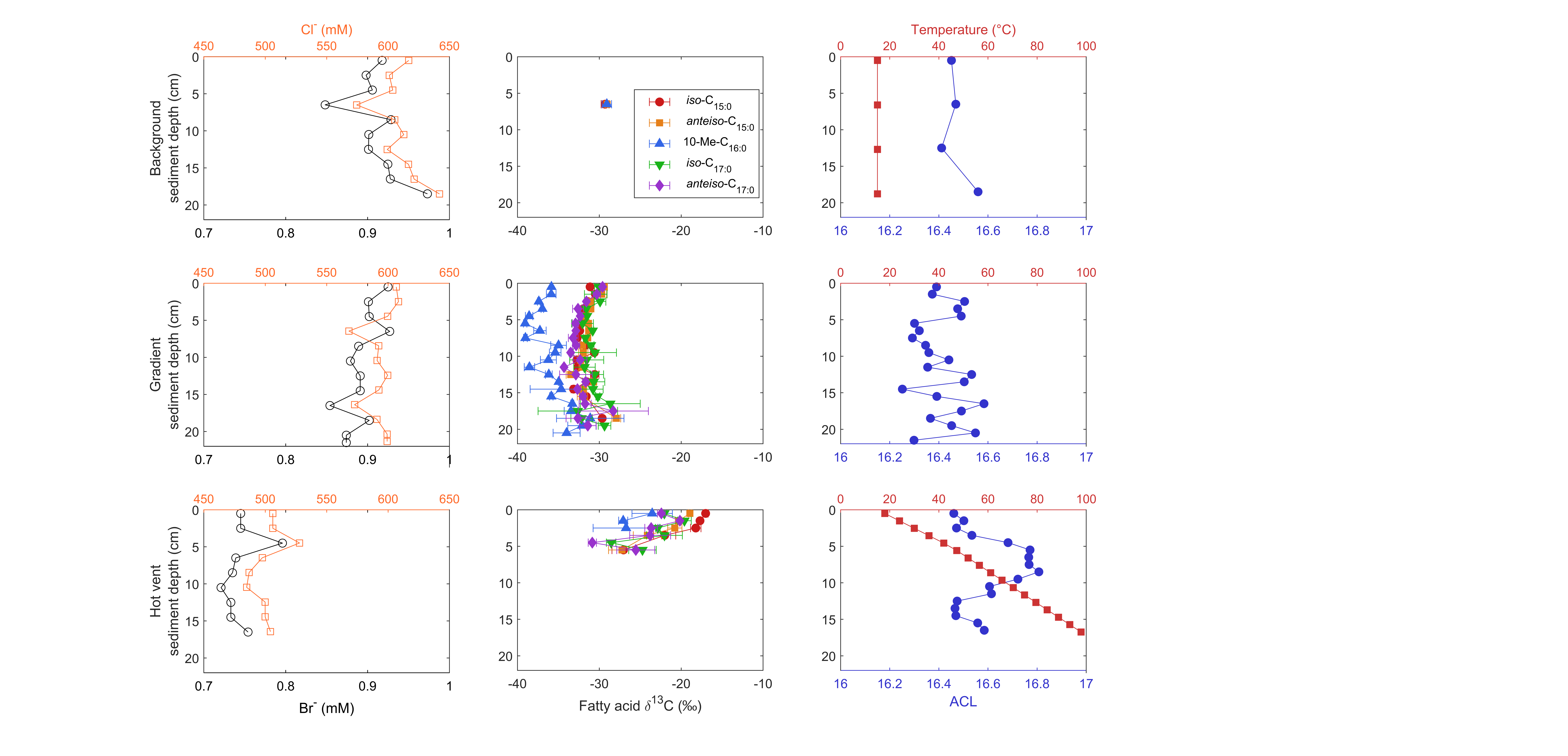

**Figure 1**: Chloride and bromide concentrations (in mM, left panel), δ^13^C values of fatty acids (in ‰, middle panel) representative of sulfate-reducing bacteria (*iso*- and *anteiso*-C_15/17_ and 10-Me-C_16:0_, e.g., Taylor and Parkes 1983, Elvert et al. 2003, Mayilraj et al. 2009), and the average chain length (ACL) of C_12_ to C_20_ fatty acids as well as *in situ* sediment temperatures measured via the temperature stick at the Background (ambient water temperature) and Hot vent site (right panel). When no δ^13^C value is depicted, fatty acid concentrations were too low for δ^13^C measurements. Temperatures from the Background were taken from measurements of bottom water using the ROV under the consideration that within 20 cm sediment depth there is no change in temperature. For the Gradient core, no temperature measurements were taken.

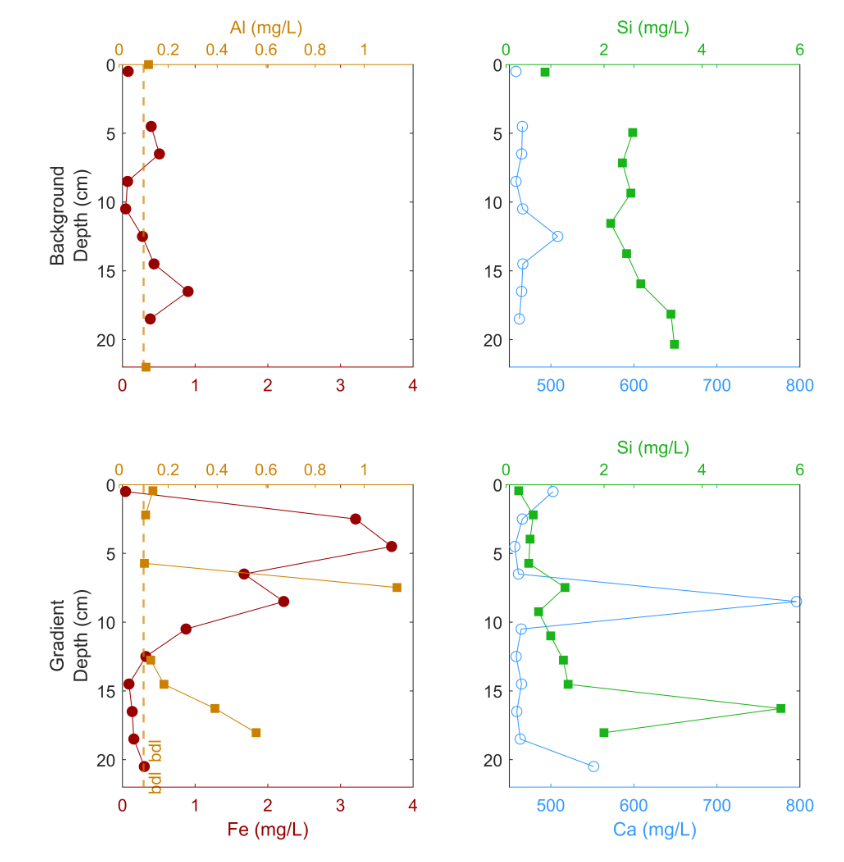

**Figure 2**: Selected total element concentrations in solution measured in the porewaters of the Background (upper panel) and Gradient core (lower panel). In the Hot vent core, no element concentrations were measured because of less porewater recovered. bdl: below detection limit.

**Table 1**: Correlation analyses using Spearmans ρ between environmental parameters (EnvVar) and fatty acid concentrations at all sites. The p-adjusted value (padj) is the p-value recalculated to control the false‐discovery rate (using Benjamini and Hochberg 1995). ACL: average chain length.

| **EnvVar** | **Fatty acid** | **Spearmans ρ** | **p.val** | **Padj** |
| --- | --- | --- | --- | --- |
| **δ^13^C DIC** | *iso*-C_17:0_ | -0.93 | 3.40E-19 | 9.85E-18 |
| **δ^13^C DIC** | *n-*C_15:0_ | -0.90 | 4.83E-16 | 7.00E-15 |
| **δ^13^C DIC** | *n-*C_17:0_ | -0.87 | 3.79E-14 | 3.66E-13 |
| **δ^13^C DIC** | *anteiso*-C_15:0_ | -0.87 | 5.45E-14 | 3.95E-13 |
| **δ^13^C DIC** | C_16:1ω7c_ | -0.84 | 1.47E-12 | 8.55E-12 |
| **δ^13^C DIC** | C_17:1ω8_ | -0.84 | 2.56E-12 | 1.24E-11 |
| **DIC** | *n-*C_15:0_ | -0.82 | 1.27E-11 | 1.84E-10 |
| **δ^13^C DIC** | C_16:1ω5_ | -0.82 | 1.74E-11 | 7.20E-11 |
| **δ^13^C DIC** | C_18:1ω7t_ | -0.82 | 2.14E-11 | 7.77E-11 |
| **pH** | δ^13^C C_16:1_ | -0.80 | 8.06E-11 | 7.79E-10 |
| **DIC** | *iso*-C_17:0_ | -0.80 | 8.91E-11 | 8.61E-10 |
| **pH** | *n*-C_12:0_ | -0.78 | 7.56E-10 | 4.39E-09 |
| **δ^13^C DIC** | *n*-C_19:0_ | -0.78 | 7.83E-10 | 2.52E-09 |
| **δ^13^C DIC** | *n*-C_18:0_ | -0.78 | 9.96E-10 | 2.89E-09 |
| **DIC** | C_17:1ω8_ | -0.78 | 1.01E-09 | 5.37E-09 |
| **DIC** | C_16:1ω7t_ | -0.77 | 1.11E-09 | 5.37E-09 |
| **DIC** | *n*-C_16:0_ | -0.77 | 1.80E-09 | 7.46E-09 |
| **Si** | *n*-C_16:0_ | -0.76 | 2.34E-09 | 6.79E-08 |
| **δ^13^C DIC** | C_16:1ω7t_ | -0.76 | 2.35E-09 | 6.19E-09 |
| **DIC** | C_18:1ω7t_ | -0.76 | 2.55E-09 | 9.24E-09 |
| **DIC** | C_16:1ω7c_ | -0.76 | 3.20E-09 | 1.03E-08 |
| **DIC** | *n*-C_17:0_ | -0.76 | 3.61E-09 | 1.05E-08 |
| **DIC** | *anteiso*-C_15:0_ | -0.76 | 3.97E-09 | 1.05E-08 |
| **Br^-^** | *n*-C_12:0_ | -0.73 | 3.08E-08 | 1.58E-07 |
| **Si** | C_18:1ω7c_ | -0.73 | 3.38E-08 | 3.35E-07 |
| **Si** | *n*-C_14:0_ | -0.73 | 3.47E-08 | 3.35E-07 |
| **δ^13^C DIC** | *n*-C_16:0_ | -0.73 | 3.74E-08 | 7.75E-08 |
| **Eh** | δ^13^C C_16:1_ | -0.72 | 5.18E-08 | 1.25E-07 |
| **Br^-^** | δ^13^C C_18:1_ | -0.72 | 5.22E-08 | 2.16E-07 |
| **δ^13^C DIC** | *iso*-C_16:0_ | -0.71 | 8.04E-08 | 1.46E-07 |
| **Eh** | δ^13^C C_18:1_ | -0.71 | 1.17E-07 | 2.62E-07 |
| **ACL** | *n*-C_15:0_ | -0.70 | 5.26E-07 | 1.19E-05 |
| **Eh** | *n*-C_12:0_ | -0.70 | 1.84E-07 | 3.81E-07 |
| **ACL** | *anteiso*-C_15:0_ | -0.69 | 8.20E-07 | 1.19E-05 |
| **DIC** | *n*-C_14:0_ | -0.69 | 3.59E-07 | 8.02E-07 |
| **Temp** | C_16:1ω5_ | -0.69 | 3.67E-07 | 1.07E-05 |
| **Br^-^** | δ^13^C C_16:1_ | -0.68 | 6.79E-07 | 1.76E-06 |
| **Si** | 10-Me-C_16:0_ | -0.67 | 9.10E-07 | 5.64E-06 |
| **ACL** | C_16:1ω7c_ | -0.67 | 2.01E-06 | 1.46E-05 |
| **Si** | C_16:1ω9_ | -0.67 | 9.72E-07 | 5.64E-06 |
| **δ^13^C DIC** | *anteiso*-C_17:0_ | -0.67 | 9.83E-07 | 1.68E-06 |
| **Temp** | *n*-C_18:0_ | -0.67 | 1.14E-06 | 1.65E-05 |
| **DIC** | C_16:1ω5_ | -0.66 | 1.23E-06 | 2.54E-06 |
| **ACL** | *n*-C_14:0_ | -0.66 | 1.29E-06 | 1.24E-05 |
| **Si** | C_16:1ω7t_ | -0.65 | 2.60E-06 | 1.08E-05 |
| **ACL** | C_17:1ω8_ | -0.64 | 6.18E-06 | 2.99E-05 |
| **ACL** | *n*-C_16:0_ | -0.64 | 4.26E-06 | 2.47E-05 |
| **Sulfide minerals** | δ^13^C C_18:1_ | -0.64 | 4.62E-06 | 6.70E-05 |
| **DIC** | *n*-C_18:0_ | -0.64 | 4.77E-06 | 9.23E-06 |
| **Cl^-^** | δ^13^C C_18:1_ | -0.63 | 5.00E-06 | 1.32E-05 |
| **SO_4_^2-^** | δ^13^C C_18:1_ | -0.63 | 5.63E-06 | 1.48E-05 |
| **ACL** | *iso*-C_15:0_ | -0.63 | 1.19E-05 | 4.92E-05 |
| **Sulfide minerals** | *n*-C_12:0_ | -0.62 | 9.83E-06 | 9.50E-05 |
| **Si** | _C18:1ω7t_ | -0.61 | 1.21E-05 | 3.79E-05 |
| **Si** | *iso*-C_15:0_ | -0.61 | 1.31E-05 | 3.79E-05 |
| **Si** | C_17:1ω8_ | -0.61 | 1.52E-05 | 3.92E-05 |
| **Cl^-^** | *n*-C_12:0_ | -0.61 | 1.54E-05 | 3.44E-05 |
| **Si** | *n*-C_15:0_ | -0.61 | 1.62E-05 | 3.92E-05 |
| **SO_4_^2-^** | *n*-C_12:0_ | -0.60 | 1.95E-05 | 4.03E-05 |
| **ACL** | C_16:1ω7t_ | -0.60 | 3.31E-05 | 0.00011999 |
| **Cl^-^** | δ^13^C C_16:1_ | -0.60 | 2.43E-05 | 5.03E-05 |
| **TOC** | *n*-C_19:0_ | 0.60 | 2.01E-05 | 0.00014559 |
| **SO_4_^2-^** | *n*-C_16:0_ | 0.61 | 1.54E-05 | 3.44E-05 |
| **Br^-^** | *n*-C_16:0_ | 0.61 | 1.27E-05 | 2.45E-05 |
| **pH** | *n*-C_18:0_ | 0.61 | 1.26E-05 | 2.82E-05 |
| **Cl^-^** | *n*-C_19:0_ | 0.61 | 1.15E-05 | 2.79E-05 |
| **Br^-^** | *n*-C_19:0_ | 0.62 | 1.13E-05 | 2.35E-05 |
| **Br^-^** | *n*-C_18:0_ | 0.62 | 1.05E-05 | 2.35E-05 |
| **SO_4_^2-^** | *n*-C_19:0_ | 0.62 | 7.52E-06 | 1.82E-05 |
| **Cl^-^** | C_16:1ω7t_ | 0.64 | 4.48E-06 | 1.30E-05 |
| **Eh** | *n*-C_16:0_ | 0.64 | 3.52E-06 | 6.00E-06 |
| **Si** | δ^13^C C_18:1_ | 0.65 | 2.03E-06 | 9.80E-06 |
| **pH** | C_16:1ω5_ | 0.66 | 1.71E-06 | 4.13E-06 |
| **sulfur** | *n*-C_14:0_ | 0.66 | 1.50E-06 | 2.17E-05 |
| **Cl^-^** | *n*-C_18:0_ | 0.67 | 1.13E-06 | 3.64E-06 |
| **Eh** | *anteiso*-C_17:0_ | 0.67 | 9.78E-07 | 1.77E-06 |
| **SO_4_^2-^** | C_16:1ω7t_ | 0.67 | 9.16E-07 | 2.66E-06 |
| **Cl^-^** | _C18:1ω7t_ | 0.67 | 7.68E-07 | 2.78E-06 |
| **Br^-^** | C_16:1ω7t_ | 0.67 | 7.26E-07 | 1.76E-06 |
| **Br^-^** | _C18:1ω7t_ | 0.68 | 6.43E-07 | 1.76E-06 |
| **Br^-^** | C_16:1ω5_ | 0.68 | 5.48E-07 | 1.76E-06 |
| **pH** | C_16:1ω7c_ | 0.68 | 5.25E-07 | 1.38E-06 |
| **pH** | C_17:1ω8_ | 0.68 | 5.12E-07 | 1.38E-06 |
| **Eh** | C_16:1ω7t_ | 0.69 | 3.07E-07 | 5.93E-07 |
| **SO_4_^2-^** | *n*-C_18:0_ | 0.69 | 2.73E-07 | 8.79E-07 |
| **Cl^-^** | C_16:1ω5_ | 0.70 | 1.56E-07 | 6.45E-07 |
| **SO_4_^2-^** | C_18:1ω7t_ | 0.71 | 1.23E-07 | 4.45E-07 |
| **pH** | *anteiso*-C_15:0_ | 0.71 | 1.13E-07 | 3.63E-07 |
| **Br^-^** | C_17:1ω8_ | 0.71 | 1.05E-07 | 3.79E-07 |
| **pH** | *iso*-C_16:0_ | 0.71 | 9.03E-08 | 3.27E-07 |
| **pH** | *n*-C_19:0_ | 0.71 | 8.70E-08 | 3.27E-07 |
| **TOC** | *anteiso*-C_17:0_ | 0.71 | 7.81E-08 | 7.55E-07 |
| **Cl^-^** | *anteiso*-C_15:0_ | 0.72 | 6.63E-08 | 3.20E-07 |
| **Cl^-^** | C_17:1ω8_ | 0.72 | 6.63E-08 | 3.20E-07 |
| **δ^13^C DIC** | *n*-C_12:0_ | 0.72 | 4.94E-08 | 9.55E-08 |
| **Cl^-^** | C_16:1ω7c_ | 0.72 | 4.13E-08 | 2.99E-07 |
| **SO4** | C_16:1ω5_ | 0.73 | 3.54E-08 | 1.47E-07 |
| **Br^-^** | C_16:1ω7c_ | 0.73 | 3.26E-08 | 1.58E-07 |
| **Br^-^** | *anteiso*-C_15:0_ | 0.73 | 2.68E-08 | 1.58E-07 |
| **TOC** | *iso*-C_16:0_ | 0.73 | 2.19E-08 | 3.17E-07 |
| **SO_4_^2-^** | *anteiso*-C_15:0_ | 0.74 | 1.96E-08 | 9.49E-08 |
| **DIC** | δ^13^C C_16:1_ | 0.74 | 1.79E-08 | 4.34E-08 |
| **Cl^-^** | *n*-C_17:0_ | 0.74 | 1.70E-08 | 1.65E-07 |
| **SO_4_^2-^** | C_17:1ω8_ | 0.74 | 1.62E-08 | 9.38E-08 |
| **Eh** | *iso*-C_16:0_ | 0.74 | 1.48E-08 | 3.91E-08 |
| **δ^13^C DIC** | δ^13^C C_18:1_ | 0.75 | 1.00E-08 | 2.24E-08 |
| **SO_4_^2-^** | *n*-C_17:0_ | 0.75 | 9.45E-09 | 6.85E-08 |
| **SO_4_^2-^** | C_16:1ω7c_ | 0.75 | 8.10E-09 | 6.85E-08 |
| **δ^13^C DIC** | δ^13^C C_16:1_ | 0.75 | 6.65E-09 | 1.61E-08 |
| **Cl^-^** | *n*-C_15:0_ | 0.75 | 6.60E-09 | 9.57E-08 |
| **SO_4_^2-^** | *n*-C_15:0_ | 0.77 | 2.08E-09 | 3.01E-08 |
| **Eh** | C_18:1ω7t_ | 0.77 | 1.71E-09 | 4.96E-09 |
| **Eh** | *n*-C_19:0_ | 0.77 | 1.62E-09 | 4.96E-09 |
| **Br^-^** | *n*-C_15:0_ | 0.77 | 1.09E-09 | 1.05E-08 |
| **DIC** | *n*-C_12:0_ | 0.78 | 9.23E-10 | 5.37E-09 |
| **Br^-^** | *n*-C_17:0_ | 0.78 | 8.83E-10 | 1.05E-08 |
| **pH** | *n*-C_15:0_ | 0.78 | 5.26E-10 | 3.81E-09 |
| **Eh** | *n*-C_18:0_ | 0.79 | 2.89E-10 | 1.05E-09 |
| **pH** | *n*-C_17:0_ | 0.81 | 7.17E-11 | 7.79E-10 |
| **Eh** | C_17:1ω8_ | 0.81 | 5.20E-11 | 2.15E-10 |
| **Cl^-^** | *iso*-C_17:0_ | 0.81 | 3.62E-11 | 1.05E-09 |
| **SO_4_^2-^** | *iso*-C_17:0_ | 0.82 | 1.42E-11 | 4.12E-10 |
| **Eh** | C_16:1ω5_ | 0.82 | 1.22E-11 | 5.90E-11 |
| **DIC** | δ^13^C C_18:1_ | 0.82 | 1.04E-11 | 1.84E-10 |
| **Eh** | C_16:1ω7c_ | 0.83 | 6.73E-12 | 3.90E-11 |
| **Br^-^** | *iso*-C_17:0_ | 0.83 | 4.24E-12 | 1.23E-10 |
| **Eh** | *anteiso*-C_15:0_ | 0.83 | 3.60E-12 | 2.61E-11 |
| **pH** | *iso*-C_17:0_ | 0.86 | 2.52E-13 | 7.31E-12 |
| **Eh** | *n*-C_17:0_ | 0.87 | 3.43E-14 | 3.31E-13 |
| **Eh** | *n*-C_15:0_ | 0.88 | 1.47E-14 | 2.12E-13 |
| **TOC** | *n*-C_20:0_ | 0.89 | 1.45E-15 | 4.20E-14 |
| **Eh** | *iso*-C_17:0_ | 0.94 | 5.60E-21 | 1.62E-19 |

**Table 2**: EDX results of point measurements (1-6) of key mineral phases from the Gradient and Hot vent core depicted in Fig. 4.

| **EDX measurements** | **Atom %** | | | | | | | | | | |
| --- | --- | --- | --- | --- | --- | --- | --- | --- | --- | --- | --- |
|  | **O** | **S** | **Fe** | **Na** | **Al** | **Si** | **Cl** | **Mg** | **K** | **Ca** | **Ti** |
| 1 | 2.69 | 60.39 | 36.92 | 0 | 0 | 0 | 0 | 0 | 0 | 0 | 0 |
| 2 | 27.32 | 7.77 | 51.44 | 1.38 | 5.03 | 4.89 | 2.16 | 0 | 0 | 0 | 0 |
| 3 | 74.49 | 0 | 0 | 0 | 12.06 | 13.45 | 0 | 0 | 0 | 0 | 0 |
| 4 | 8.1 | 59.32 | 31.27 | 0 | 0.5 | 0.53 | 0 | 0.28 | 0 | 0 | 0 |
| 5 | 65.41 | 1.79 | 3.27 | 0.47 | 3.25 | 23.82 | 0.11 | 0.74 | 1.14 | 0 | 0 |
| 6 | 73.28 | 0.18 | 0.61 | 0.4 | 2.01 | 3.66 | 0 | 0.48 | 0.12 | 0.06 | 19.2 |

**References**

Benjamini, Y. & Hochberg, Y. (1995) J. R. Stat. Soc., Ser. B, Stat. Methodol. , 57:289-300. <https://doi.org/10.1111/j.2517-6161.1995.tb02031.x>

Elvert, M., Boetius, A., Knittel, K. & Jørgensen, B. B. (2003) Geomicrobiol. J., 20:403-419. <https://doi.org/10.1080/01490450303894>

Mayilraj, S., Kaksonen, A. H., Cord-Ruwisch, R., Schumann, P., Spröer, C., Tindall, B. J. & Spring, S. (2009) Extremophiles*,* 13:247-255. <https://doi.org/10.1007/s00792-008-0212-4>

Taylor, J. & Parkes, R. J. (1983) Microbiology*,* 129:3303-3309. <https://doi.org/10.1099/00221287-129-11-3303>
